# Candidates for Drug Repurposing to Address the Cognitive Symptoms in Schizophrenia

**DOI:** 10.1101/2022.03.07.483231

**Authors:** Elise Koch, Karolina Kauppi, Chi-Hua Chen

## Abstract

In the protein-protein interactome, we have previously identified a significant overlap between schizophrenia risk genes and genes associated with cognitive performance. Here, we further studied this overlap to identify potential candidate drugs for repurposing to treat the cognitive symptoms in schizophrenia. We first defined a cognition-related schizophrenia interactome from network propagation analyses, and identified drugs known to target more than one protein within this network. Thereafter, we used gene expression data to further select drugs that could counteract schizophrenia-associated gene expression perturbations. Additionally, we stratified these analyses by sex to identify sex-specific pharmacological treatment options for the cognitive symptoms in schizophrenia. After excluding drugs contraindicated in schizophrenia, we identified eight drug candidates, most of which have anti-inflammatory and neuroprotective effects. Due to gene expression differences in male and female patients, four of those drugs were also selected in our male-specific analyses, and the other four in the female-specific analyses. Based on our bioinformatics analyses of disease genetics, we suggest eight candidate drugs that warrant further examination for repurposing to treat the cognitive symptoms in schizophrenia, and suggest that these symptoms could be addressed by sex-specific pharmacological treatment options.

## 1. Introduction

Cognitive symptoms are considered as a core feature of schizophrenia, affecting about 80 % of the patients [1,2]. Impairments within various cognitive domains are observed already in the premorbid phases of schizophrenia [3,4]. While these cognitive symptoms have a higher impact on functional outcomes in schizophrenia than the positive symptoms [5–7], currently available medications used in schizophrenia primarily target the positive symptoms by decreasing dopamine in the brain [8]. Thus, cognitive symptoms constitute an important unmet treatment need for individuals with schizophrenia or other psychiatric disorders. Despite attempts to develop drugs that address the cognitive symptoms in schizophrenia or to investigate existing drugs with potential procognitive effects, no significant clinical progress has been made so far, and there are currently no available medications that efficiently treat the cognitive symptoms in psychiatric disorders [9,10].

Drugs which target, e.g. a receptor or an enzyme encoded by a gene in which genetic variants associate with the target disease have shown a higher success rate in the drug development pipeline [11]. Network medicine is an approach in which gene networks are defined through protein-protein interactions (PPIs) of gene products. The important concept that the network approach embodies is that the effect of a mutation in one gene may impact its neighbors in the network of PPIs. Therefore, it is important to take PPIs into account in the effort to reveal genetic disease mechanisms and to identify drug targets [12,13]. To identify existing drugs that potentially can be used for repurposing to treat conditions other than their original indication, interactions of protein products from disease risk genes can be studied within gene networks [14]. This is important considering the fact that the number of new treatments coming to the market remains low [15], and drug repurposing is a solution with reduced costs and safety concerns [14]. More than 31% of GWAS-associated SNPs are pleiotropic [16], which provides an explanation why several drugs have been successfully repurposed. One of the reasons why genes are pleiotropic is that gene products are connected to each other by different mechanisms such as protein-protein interactions, thus affecting different biological pathways that can affect several clinical outcomes [17]. Therefore, network-based drug-disease proximity within networks of protein-protein interactions can elucidate the relationship between drugs and diseases, and serves as a useful tool to identify new indications for approved drugs with known safety profiles [18]. However, network proximity is not sufficient for a drug to be effective, as drugs also need to induce the right perturbation in the cell [19].

Complex diseases such as schizophrenia are considered to be partly driven by differentially expressed genes (DEG) [20], and drugs administered to treat these diseases often revert the DEG to their normal levels [21]. To prioritize repurposed drug candidates based on network proximity, gene expression profiles for both the disease and the candidate drugs can be compared to select drug candidates that may counteract disease-induced gene expression perturbations (by down-regulating genes up-regulated in the disease or vice versa), and exclude drugs that may enhance disease-related perturbations. Improved knowledge of cognition-related schizophrenia genetics in the context of gene networks incorporating gene expression provides a platform to find existing drugs for repurposing candidates to treat the cognitive symptoms in schizophrenia [21,22].

Within biological gene networks, we have previously identified a significant interactome overlap between schizophrenia risk genes and genes associated with cognitive performance, and identified a cognition-related subset of schizophrenia risk genes [23]. Here, we first redefined cognition-related schizophrenia risk genes in the updated human protein interactome [24]. Then, we further studied these genes and their network neighbors in relation to interactions with existing drugs, to identify potential repurposing candidates. Thereafter, we selected drug candidates with opposite direction of effect on gene expression in brain tissues than that observed in schizophrenia patients relative to controls, for suggestion as candidates for drug repurposing to identify whether any of these drugs may potentially be repurposable to effectively treat the cognitive symptoms in schizophrenia. However, gene expression perturbations related to schizophrenia differ between males and females [25]. Therefore, we also retrieved gene expression data separately for males and females, and performed sex-stratified analyses to identify drug repurposing candidates with potential for sex-specific effects on cognitive symptoms in schizophrenia.

## 2. Materials and Methods

### 2.1. Disease/trait Genes

To identify genes associated with schizophrenia and cognitive functioning, summary statistics from large-scale GWASs were used. For schizophrenia, we used a GWAS performed by the Psychiatric Genomics Consortium (PGC) including 36,989 cases and 113,075 controls, which identified 108 independent loci associated with schizophrenia [26]. For cognitive functioning, we used a large multicenter GWAS on cognitive performance measured across at least three domains of cognitive performance including 257,841 individuals [27]. For both phenotypes, genes linked to genome-wide significant single nucleotide polymorphisms (SNPs) were identified using the “SNP2GENE” function of the web-based software Functional Mapping and Annotation of Genome-Wide Association Studies (FUMA) [28], after excluding the major histocompatibility complex (MHC) region (25-34 Mb on chromosome 6 on the hg19 assembly) due to the high linkage disequilibrium (LD) of this region. Genes were selected from genomewide significant SNPs (p < 5 x 10^-8^) that were located within a LD-threshold of r^2^ < 0.6, and a maximum distance of 250 kb, with a minor allele frequency (MAF) of ≥ 0.01 (positional mapping with default settings in FUMA). For schizophrenia, the genes C4A and C4B were included to represent the signal from the MHC region, because it has been shown that SNPs incorporating these genes are risk factors for schizophrenia [29,30].

### 2.2. The Human Protein Interactome

The human interactome [12,24] was constructed from data of 15 commonly used databases, focusing on high-quality protein-protein interactions (PPIs) as follows: Physical PPIs tested by high-throughput yeast-two-hybrid (Y2H) screening system; literature-curated PPIs followed by affinity-purification mass spectrometry (AP-MS), Y2H, and literature-derived low-throughput experiments; physical PPIs derived from protein three-dimensional structures; kinase-substrate interactions by literature-derived low-throughput and high-throughput experiments; and signaling networks by literature-derived low-throughput experiments. For details see Cheng et al. 2019 [24]. The human interactome was updated on 2021-08-11, and consists of 16,706 proteins interconnected by 246,995 interactions. Network figures were created using Cytoscape [31], where nodes refer to genes or drugs, and edges refer to gene-drug interactions or genegene interactions through identified PPIs between gene products (proteins).

### 2.3. Cognition-related Schizophrenia Interactome

To define a cognition-related schizophrenia interactome, we used the method network propagation [32–34], implemented in the Cytoscape application Diffusion [32]. Starting with a chosen set of input proteins, information from their PPIs is transferred to all other proteins in the network and received from them through an iterative process. Network proximity between proteins is scored depending on their PPIs, where higher diffusion output values relate to higher relatedness to the input proteins [32–34].

First, we performed a network propagation analysis with all cognition-associated genes as input query to define which schizophrenia risk genes are close to genes associated with cognitive performance. This was done in the updated human interactome where we excluded self-interactions, resulting in 243,603 interactions connecting 16,706 proteins. To define which genes are close to cognition-associated genes, we calculated the Youden index [35] to determine a cut-off from 1,000 ROC analyses with the diffusion output values from all cognition-associated genes and the diffusion output values from non-cognition random genes with equal number and comparable node degree [13] as the cognition-associated genes. Two-sample proportion tests were performed to test if schizophrenia risk genes are significantly overrepresented among genes defined as close to cognition. To investigate if SNPs related to schizophrenia risk genes defined as close to cognition have lower p-values in the GWAS of cognitive performance compared to schizophrenia risk genes not defined as close to cognition, we performed a Welch two sample t-test. SNPs were positionally mapped to genes adding 25 kb upstream and downstream.

As most approved drugs do not target disease proteins, but bind to proteins in their network vicinity [36], we next defined a cognition-related schizophrenia interactome including not only the schizophrenia risk genes that were defined as close to cognition-associated genes, but also genes in their immediate network proximity. The schizophrenia risk genes that were defined as close to cognition-associated genes were used as input query genes in another network propagation analysis, and a cut-off was determined by the Youden index from 1,000 receiver operating characteristic (ROC) analyses with the diffusion output values from the input genes and the diffusion output values from non-input genes with equal number and comparable node degree as the input genes to define a cognition-related schizophrenia interactome. ROC analyses were performed using the pROC package [37] in R version 4.0.3.

### 2.4. Drug Target Network

The drug-gene interaction database (DGIdb) [38] (version v4.2.0, last updated 2021-04-13) was used to identify drug-gene interactions between approved drugs and genes in the cognition-related schizophrenia interactome. The DGIdb provides information on drug-gene interactions from 22 sources that are aggregated and normalized (for description of sources in DGIdb, see Supplementary Table 1 in Freshour et al. 2021 [38]).

### 2.5. Gene Expression Perturbation Profiles

For the drugs interacting with genes in the cognition-related schizophrenia interactome, we utilized gene expression data (drug versus no drug) to evaluate if these drugs modulate the activity of the genes in our network. To determine each drug’s gene expression perturbation profile, we retrieved gene expression data from the Connectivity Map (CMAP) database [39,40], extracted from the Phase 2 data release of the Library of Integrated Cellular Signatures (LINCS) in GEO series GSE70138 (GSE70138_Broad_LINCS_Level5_COMPZ_n118050×12328_2017-03-06.gctx.gz available at https://www.ncbi.nlm.nih.gov/geo/query/acc.cgi?acc=GSE70138) using cmapR package [41] in R version 4.0.3.

Gene expression data from the Gene Expression Omnibus (GEO) database [42] was used to evaluate schizophrenia-induced gene expression perturbations. Data on gene expression (expression profiling by array) in different brain regions of postmortem brains from schizophrenia cases versus controls were retrieved from eight GEO datasets (available at https://www.ncbi.nlm.nih.gov/geo/): GSE53987, GSE21935, GSE17612, GSE12654, GSE62191, GSE35978, GSE21138, GSE12649 (comprising 226 cases and 219 controls in total). DEGs between schizophrenia cases and controls were identified using GEO2R (https://www.ncbi.nlm.nih.gov/geo/geo2r/), an online tool to identify DEGs from GEO series [42]. To combine the outputs from the eight datasets, the following steps were done. For each gene within the cognition-related schizophrenia interactome, we first calculated the mean fold change (FC) for each of the eight series. Then, we z-transformed these mean FC values of all selected genes within each study, to generate values that are comparable across the eight studies. Thereafter, we calculated the mean of the z-transformed mean FC values across the eight studies to retrieve one FC value per gene in our network.

To evaluate if the drug candidates could change schizophrenia-induced gene expression perturbations (whether they down-regulate genes up-regulated in schizophrenia or vice versa), we calculated the Spearman correlation ρ between the drug-induced perturbations and the schizophrenia-induced perturbations in genes within the cognition-related schizophrenia interactome. This was done for each drug in the cognition-related schizophrenia interactome, where negative correlation coefficients indicate that the drug could counteract schizophrenia-associated effects. Then, only drugs with negative correlation coefficients (regardless of correlation p-values) were selected for further consideration as repurposing candidates, thereby excluding drugs that may worsen disease-related gene-expressions perturbations.

In addition, we evaluated schizophrenia-induced gene expression perturbations separately for males and females. Of the eight GEO datasets used, data on sex was available for five series: GSE53987, GSE21935, GSE17612, GSE35978, GSE21138. In total, these series comprise 149 schizophrenia cases (102 males and 47 females) and 140 controls (91 males and 49 females). DEGs between schizophrenia cases and controls were retrieved separately for males and females, and the mean FC per gene was calculated as described above. For each gene within the cognition-related schizophrenia interactome, the correlation between these sex-specific FC values and the CMAP-retrieved drug perturbations was calculated separately for males and females.

### 2.6. Prioritization of Repurposing Candidates

From the drugs interacting with proteins in our cognition-related schizophrenia interactome, we first excluded drugs interacting only with genes coding for the drug-metabolizing enzymes of the Cytochrome P450 (CYP) family present in our network (CYP2D6 and CYP2C9), as many drugs are metabolized by these enzymes [43]. Thereafter, we further investigated those drugs whose gene expression perturbation profile was negatively correlated with schizophrenia-induced gene expression perturbations. From those, we excluded drugs interacting with only one protein in our network, as network pharmacology analyses have revealed that drugs acting on a single drug target within a disease network are often not effective [44,45]. Furthermore, we excluded antipsychotics and drugs with severe side effects or contraindications to schizophrenia.

## 3. Results

From the schizophrenia GWAS summary statistics [26], 313 schizophrenia risk genes were identified using FUMA [28], and 2 genes (C4A and C4B) were added to represent the signal from the MHC region, resulting in 315 schizophrenia risk genes of which 282 were present in the human interactome (**Supplementary Table 1**). For cognitive performance, 621 associated genes were identified from the corresponding GWAS summary statistics [27]. Of these, 566 were present in the human interactome (**Supplementary Table 1**).

### 3.1. Cognition-related Schizophrenia Interactome

To identify specific schizophrenia risk genes that are close to cognition-associated genes, we performed a network propagation analysis with all cognition-associated genes as input query, and determined a cutoff (0.0395339) to define a subset of genes in the whole interactome that are close to cognition (N = 1,805). Schizophrenia risk genes were significantly overrepresented among all genes defined as close to cognition (N = 72, z-value = 8.035, p-value < 2.2e-16), even when we excluded genes that are both schizophrenia and cognition-associated (N = 58, z-value = 5.467, p-value < 2.2e-16). In the GWAS summary statistics for cognitive performance, SNPs related to schizophrenia risk genes defined as close to cognition had significantly lower p-values than SNPs related to schizophrenia risk genes not defined as close to cognition (t-value = 31.994, p-value <2.2e-16), even when we excluded genes that are both schizophrenia and cognition-associated (t-value = −2.560, p-value = 0.0106). The specific schizophrenia risk genes defined as close to cognition (N = 72) can be found in **Supplementary Table 1**. Using these genes as input query genes in another network propagation analysis, a cut-off (0.0057976) was determined to define a cognition-related schizophrenia interactome including 735 genes of which 76 are GWAS-significant schizophrenia risk genes and 104 are GWAS-significant cognition genes (listed in **Supplementary Table 1**).

### 3.2. Drug Repurposing Candidates for the Cognitive Symptoms in Schizophrenia

From the DGIdb [38], drug-gene interactions were identified between 751 approved drugs and 155 genes in the cognition-related schizophrenia interactome. After exclusion of drugs that only interact with genes coding for CYP450 enzymes (N = 236) as well as drugs not present in the CMAP (N = 290), 225 drugs interacting with 101 proteins in the network were further investigated (**Figure 1**). Schizophrenia-associated gene expression perturbations were available for 91 of the 101 genes (**Supplementary Table 2**). Out of the 225 drugs, 47 showed opposite gene expression perturbations in drug (drug-induced expression) versus schizophrenia (disease-associated expression) in the 101 corresponding genes (**Figure 2, Supplementary Table 3**). From these 47 drugs, we excluded drugs interacting with only one gene in our network (N = 20), antipsychotics (N = 5), as well as drugs with severe side effects or contraindications to schizophrenia such as drugs increasing dopamine potentially worsening positive symptoms (N = 14), resulting in eight drug candidates (**Figure 1, Figure 3A**). The eight drug candidates, their indications, as well as their interaction partner in the cognition-related schizophrenia interactome are listed in **Table 1**. Information on indications was collected from the DrugBank (https://go.drugbank.com) [46].

**Figure 1.**
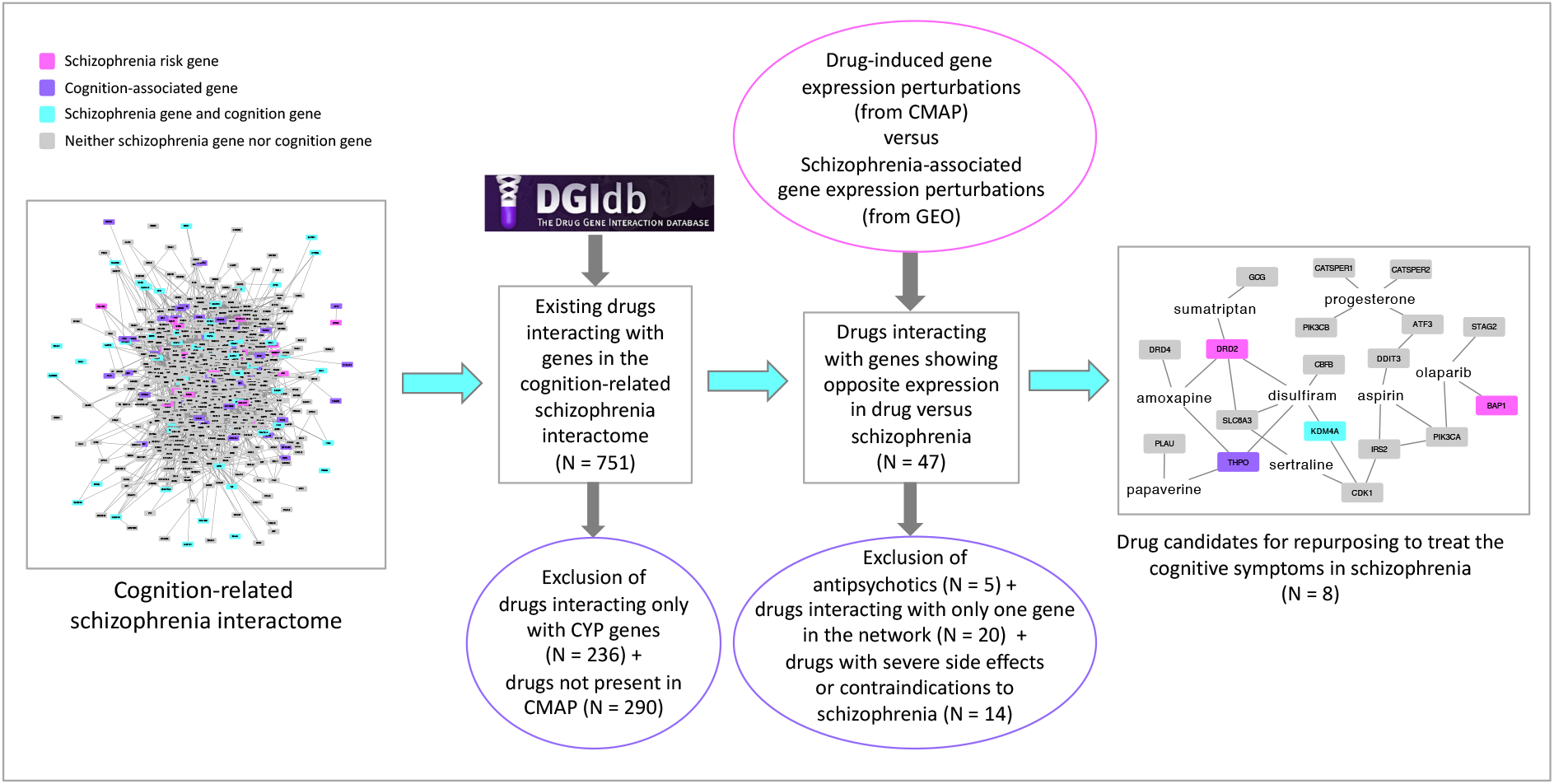
Flow chart depicting the steps taken in this study to evaluate drugs interacting with proteins in the cognition-related schizophrenia interactome to prioritize the shortlisted drugs. N = number of drugs

**Figure 2.**
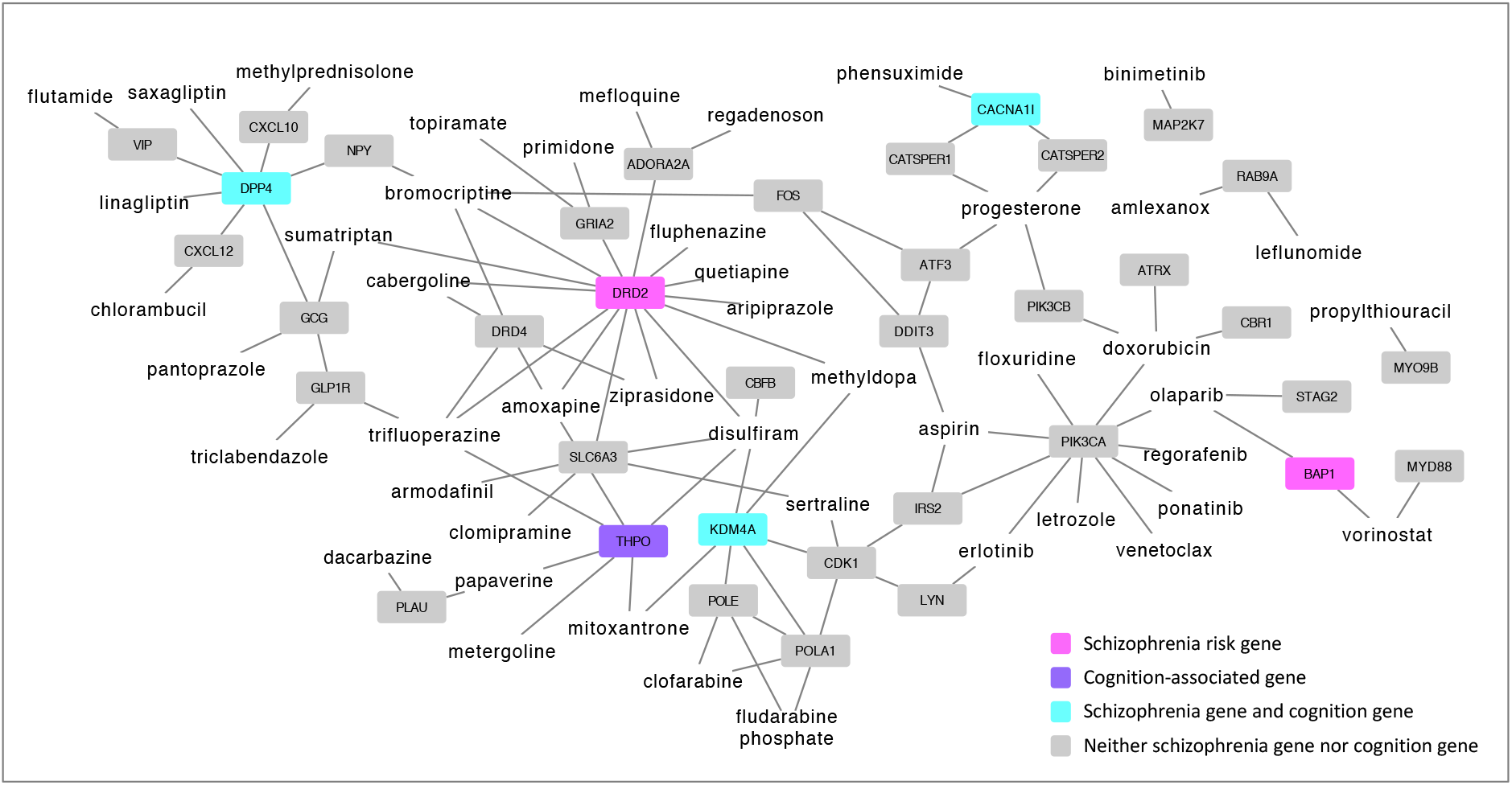
Drugs whose gene expression perturbation profile was negatively correlated with schizophrenia-associated gene expression perturbations across 101 genes in the cognition-related schizophrenia interactome (N = 47 drugs), shown with their protein interaction partners in the network (N = 37 proteins), excluding CYP2D6 and CYP2C9.

**Figure 3.**
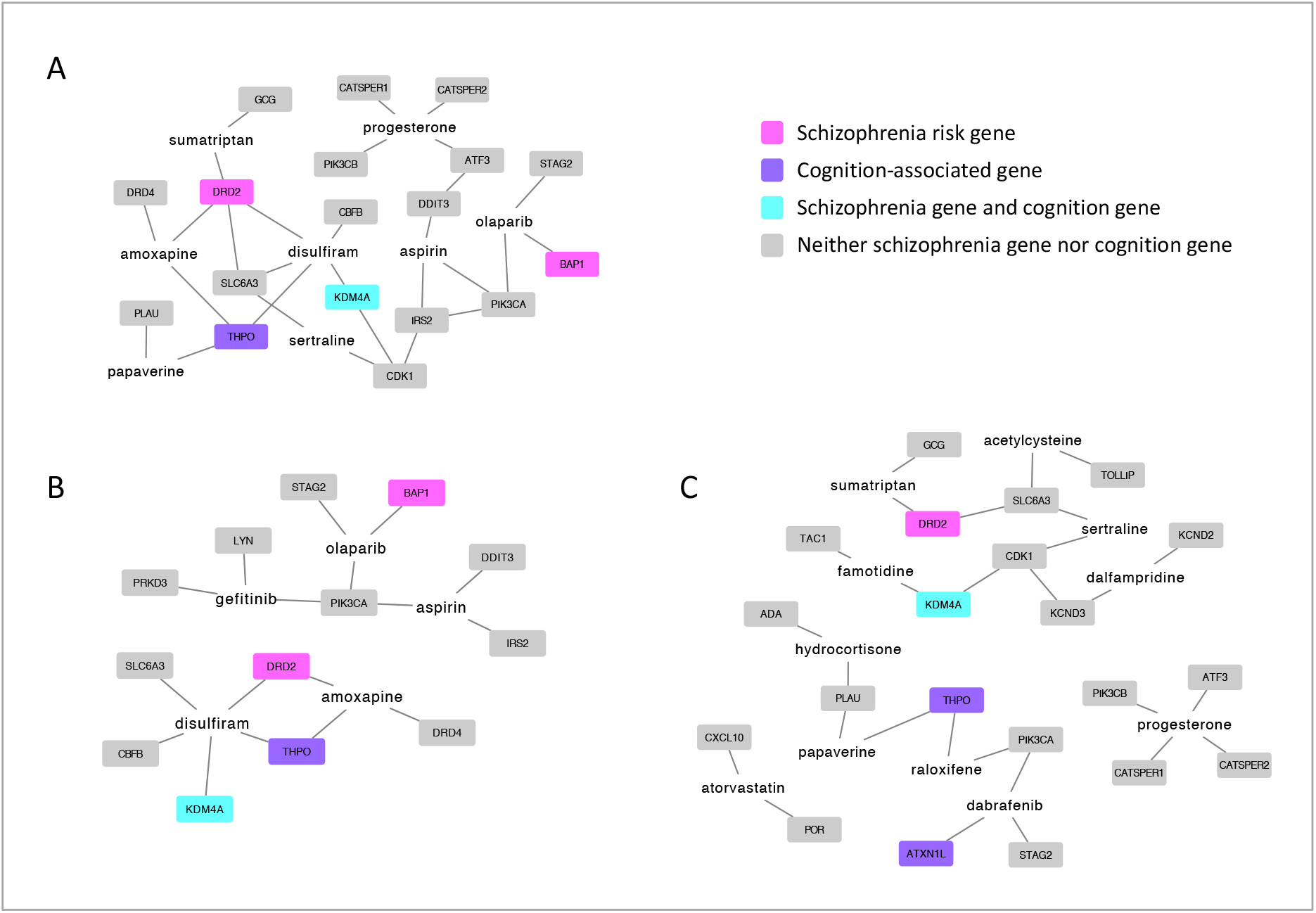
Drug candidates for repurposing to address the cognitive symptoms in schizophrenia (N = 8) based on drug-induced versus schizophrenia-induced gene expression in all individuals (**A**), as well as male-specific (N = 5) (**B**) and female-specific (N = 11) (**C**) repurposing candidates based on drug-induced versus sex-specific schizophrenia-induced gene expression, and their interaction partners in the cognition-related schizophrenia interactome, excluding CYP2D6 and CYP2C9.

**Table 1:** Drug candidates for repurposing to treat the cognitive symptoms in schizophrenia.

| **Drug** | **Interaction partners in the network** | **Indication** |
| --- | --- | --- |
| Amoxapine | DRD2, DRD4, THPO, CYP2D6 | Neurotic or reactive depressive disorders and endogenous or psychotic depression |
| Aspirin | DDIT3, IRS2, PIK3CA, CYP2C9, CYP2D6 | Pain, fever, and inflammation |
| Disulfiram | CBFB, DRD2, KDM4A, SLC6A3, THPO, CYP2C9 | Alcohol addiction |
| Olaparib | BAP1, PIK3CA, STAG2 | Recurrent or advanced ovarian cancer and metastatic breast cancer |
| Papaverine | PLAU, THPO, CYP2C9, CYP2D6 | Smooth muscle spasms |
| Progesterone | ATF3, CATSPER1, CATSPER2, PIK3CB, CYP2D6, CYP2C9 | Contraception, control of abnormal uterine bleeding, prevention of endometrial hyperplasia |
| Sertraline | CDK1, SLC6A3 | Major depressive disorder, social anxiety disorder and many other psychiatric conditions |
| Sumatriptan | DRD2, GCG | Migraine |

### 3.3. Sex-specific Drug Repurposing Candidates for the Cognitive Symptoms in Schizophrenia

To identify sex-specific drug repurposing candidates, DEGs between schizophrenia cases and controls were retrieved separately for males and females, and the correlation between sexspecific FC values and the CMAP-retrieved drug perturbations were calculated separately for males and females. Evaluating drug candidates in the same way as before, we retrieved 11 drug candidates for females and 5 drug candidates for males (**Figure 3B and C**). Out of the eight drug candidates that were identified for all individuals (**Figure 3A**), four were identified in males only (amoxapine, aspirin, disulfiram, olaparib), and the other four drugs were identified in females only (papaverine, progesterone, sertraline, sumatriptan) (**Figure 3B and C**). For males, one additional drug was identified, the anti-cancer drug gefitinib (**Figure 3B**), and for females, seven additional drugs were identified as follows: Acetylcysteine (used in chronic bronchitis), atorvastatin (used to lower abnormal cholesterol and lipid levels), dabrafenib (anti-cancer drug), dalfampridine (used in Multiple Sclerosis), famotidine (used to treat ulcer diseases), hydrocortisone (used to treat dermatoses, endocrine disorders, immune conditions, and allergic disorders), and raloxifene (used to treat osteoporosis and to prevent invasive breast cancer in high-risk postmenopausal women) (**Figure 3C**). Correlation coefficients for all 225 drugs interacting with 101 proteins in the network can be found in **Supplementary Table 3** separately for males and females.

## 4. Discussion

In this study, we have identified eight existing drugs that could potentially be used for repurposing to address the cognitive symptoms in schizophrenia. First, we defined a cognition-related schizophrenia interactome via network-based methods, and studied the genes in this network in relation to interactions with existing drugs. Then, we selected drug candidates that could change schizophrenia-induced gene expression perturbations and identified eight drugs that may potentially be repurposable to address the cognitive symptoms in schizophrenia. In addition, we evaluated drug-induced versus schizophrenia-associated gene expression profiles stratified by sex, and identified different repurposing candidates for male and female patients with schizophrenia.

Some of the eight drugs have previously been suggested for the treatment of cognitive symptoms in neuropsychiatric disorders. Aspirin, a non-steroidal anti-inflammatory drug (NSAID), has been suggested both for the prevention of Alzheimer’s disease (AD) [47] and for the treatment of schizophrenia [48], due to the emerging evidence of excessive inflammatory response in the pathophysiology of both schizophrenia [48] and AD [47]. For the prevention of dementia, evidence is insufficient to fully evaluate the effect of aspirin on cognitive decline and risk of dementia [49,50]. Results from clinical trials investigating effects of adding aspirin to standard antipsychotic treatment in schizophrenia are inconclusive [51,52]. Although some improvements of positive and negative symptoms have been reported, cognitive measures were not included in these studies [51,52]. This suggests the need for future studies investigating cognitive effects of aspirin in schizophrenia.

Papaverine is an inhibitor of phosphodiesterase (PDE) 10A, an enzyme that catalyzes the hydrolysis of the second messenger molecules cyclic-3’,5’-adenosine monophosphate (cAMP) and cyclic-3’,5’-guanosine monophosphate (cGMP), thereby controlling cyclic nucleotide signaling and cellular communication [53,54]. As both cAMP and cGMP are involved in cognition [55], neuroprotection [56,57] and inflammation [58,59], and PDEs are highly expressed in the immune and central nervous system (CNS), papaverine and other PDE10A inhibitors have emerged as promising drugs for the treatment of CNS disorders [60]. Both antipsychotic and procognitive effects have been demonstrated in preclinical studies on PDE10A inhibitors for the treatment of schizophrenia [54,61], but clinical trials investigating different PDE10A inhibitors for the treatment of schizophrenia have not been successful so far [61]. However, it has recently been shown that papaverine has anti-inflammatory and neuroprotective effects [62,63] suggesting that papaverine has the therapeutic potential for improving cognitive dysfunctions in neuropsychiatric and neurodegenerative disorders.

Two antidepressant drugs were identified, amoxapine and sertraline. The tricyclic antidepressant (TCA) amoxapine has shown efficacy as an antipsychotic in schizophrenia [64–66]. Whereas it has been shown that amoxapine has similar effects on positive and negative symptoms as commonly used antipsychotics, its cognitive effects in schizophrenia have not been evaluated [64–66]. Interestingly, amoxapine has been identified to reduce amyloid-β production, suggesting that amoxapine could improve cognitive function and potentially prevent AD [67]. Studies investigating cognitive effects of antidepressants have demonstrated procognitive effects in depression, but findings between different agents are mixed [68], with selective serotonin reuptake inhibitors (SSRIs) showing the greatest procognitive effects [69]. The SSRI sertraline has demonstrated neuroprotective and procognitive effects in vitro and in vivo [70]. In a study of AD patients, it has been found that sertraline treatment was associated with superior effectiveness in relation to cognitive symptoms compared to treatment with the selective norepinephrine reuptake inhibitor (SNRI) venlafaxine and the TCA desipramine [71]. However, whether amoxapine or sertraline have procognitive effects in schizophrenia has to be evaluated in future studies.

Sumatriptan is a 5HT_1B/D_ receptor agonist commonly used to treat migraine attacks [72]. The number of studies examining sumatriptan’s CNS effects is low, because it has been long assumed that sumatriptan and other triptans do not penetrate the CNS [73]. In individuals with migraine, it has been shown that sumatriptan restores migraine-related drops in cognitive function during migraine attacks [74–76]. In a study of healthy females receiving sumatriptan, it has been shown that sumatriptan caused a significant decrease in reaction time for recognition of words compared to placebo, suggesting CNS effects of sumatriptan [77]. In addition to its antimigraine properties via activation of serotonin 5HT_1B/D_ autoreceptors, sumatriptan has been recognized as an anti-inflammatory agent [78]. Current evidence from preclinical studies suggests that low doses of sumatriptan are useful in many inflammatory conditions, which has to be further investigated in clinical studies [78].

Disulfiram has been widely used to treat alcoholism for over 60 years without severe side effects [79]. Cognitive effects of disulfiram in non-alcohol-dependent individuals have been evaluated in one study showing no effect on cognition after two weeks of disulfiram treatment [80]. However, no studies on cognitive effects of long term disulfiram treatment exist [81]. It has recently been shown that disulfiram prevents pyroptosis and release of the pro-inflammatory cytokine interleukin 1β, suggesting disulfiram as a repurposing candidate to counteract inflammation in various diseases [82]. In addition, disulfiram has recently been identified as a repurposing candidate to treat various cancers [83].

In network-based analyses, four anti-cancer drugs have recently been identified as potential repurposing candidates to treat AD, probably because these drugs ameliorate AD-associated neurotoxicity and neuroinflammation [84]. One of these drugs was gefitinib that we identified as a repurposing candidate in males. We also identified the anti-cancer drug olaparib, an inhibitor of the nuclear DNA binding enzyme poly ADP-ribose polymerase-1 that has an important role in both DNA repair and inflammatory diseases, suggesting anti-inflammatory and potentially precognitive effects of olaparib [85]. In fact, olaparib has recently been shown to downregulate neuroinflammation and improve cognitive function in mice [86]. The anticancer drug dabrafenib that we identified in females, also has anti-inflammatory effects [87].

Sex differences have long been observed in schizophrenia patients [88] with females being less affected by the disease than males [89,90]. This has led to the estrogen hypothesis postulating that estrogen exerts a protective role in the pathophysiology of schizophrenia [91]. It has also been suggested that estrogens may help alleviate some of the cognitive symptoms of schizophrenia in women, most likely because estrogens have anti-inflammatory effects by blocking the effects of cytokines [92]. The other female sex hormone, progesterone, may have similar protective effects as estrogen [93], and has been suggested to be a key modulator of central systems implicated in both the psychotic and cognitive facets of schizophrenia, but the exact actions of progesterone in schizophrenia are not known [93]. Although progesterone levels have been positively associated with cognitive performance [94], future studies are needed to evaluate progesterone as a potential agent to treat the cognitive symptoms in female schizophrenia patients.

Additional drugs that were identified in females only included two drugs used to treat inflammatory conditions, hydrocortisone and famotidine, of which famotine already had shown effectiveness in reducing positive schizophrenia symptoms [95], but its potential effects on cognitive symptoms in schizophrenia have not been studied. Moreover, dalfampridine has shown procognitive effects in patients with multiple sclerosis [96], and acetylcysteine has already been suggested to be effective for cognitive symptoms in schizophrenia [97]. The potential procognitive effects of acetylcysteine in schizophrenia may probably be mediated by various anti-inflammatory and neuroprotective processes [97]. Also, atorvastatin and raloxifene have previously been suggested for repurposing to treat schizophrenia. Atorvastatin has demonstrated some effectiveness in reducing negative symptoms in schizophrenia [98]. Atorvastatin might improve learning and memory by inhibiting inflammatory responses in the progression of AD [99], and the use of atorvastatin has been associated with somewhat lower incidence of dementia [100] as well as reduced neuroinflammation and delayed cognitive decline in atrial fibrillation patients [101]. The selective estrogen receptor modulator raloxifene in combination with antipsychotics has shown beneficial effects on positive, negative, and cognitive symptoms in both women and men with schizophrenia [92,102,103].

We have previously shown that polygenic risk for schizophrenia has sex-specific effects on cognitive performance [104] and related brain activity [105] in healthy individuals, indicating that the effect of underlying schizophrenia genetics on cognition is dependent on biological processes that differ between the sexes, suggesting that the cognitive symptoms in schizophrenia should be addressed by sex-specific pharmacological treatment options. Moreover, it has been demonstrated that, although male and female schizophrenia patients share many of the common pathways involving the same proteins, sex appears to be a major determinant of gene expression changes related to schizophrenia, with female patients showing larger gene expression changes relative to healthy individuals than male patients do [25]. Here, we evaluated drug-induced versus schizophrenia-associated gene expression profiles stratified by sex, and identified different repurposing candidates for male and female schizophrenia patients. Taken together, these results as well as known sex differences in schizophrenia [102,106,107] strongly suggest that pharmacological schizophrenia treatments should be sexspecific due to different pathophysiology between males and females.

However, the correlation coefficients shortlisting drugs whose gene expression perturbation profile was negatively correlated with schizophrenia-induced gene expression perturbations were quite low with only small differences between males and females, and therefore it could be a power issue that four drugs were identified in males only and the other four were identified in females only. It should also be noted that it is not known if the shortlisted drugs effectively counteract schizophrenia-associated gene expression. Gene expression analyses were performed to exclude drugs that potentially could worsen schizophrenia symptoms. The drug-induced gene expression profiles were based on experiments in cancer cell lines. However, such data have been shown to be of value for repurposing drugs even for non-cancer diseases [108]. Based on negative correlations of drug-induced versus disease-associated gene expression profiles, Topiramate, an anticonvulsant drug used in the treatment of epilepsy, was identified to be potentially repurposable for inflammatory bowel disease (IBD), which has been validated in a rodent model [108]. Here, we combine network-based analyses with gene expression profiles and shortlist eight drugs that potentially could be used for repurposing to treat the cognitive symptoms in schizophrenia. These results require follow-up experiments and finally clinical trials to enable clinical translation.

## 5. Conclusions

Our results pinpoint eight candidate drugs for clinical translation that potentially could be used for repurposing to treat the cognitive symptoms in schizophrenia, and suggest that the cognitive symptoms in schizophrenia could be addressed by sex-specific pharmacological treatment options.

## Author Contributions

Conceptualization: C-H.C., K.K., and E.K.; methodology: E.K., C-H.C. and K.K.; analyses: E.K.; investigation: E.K.; draft preparation: E.K.; reviewing and editing: E.K., K.K. and C-H. C.; visualization: E.K.; funding acquisition: K.K., and C-H.C. All authors read and approved the final manuscript.

## Data Availability Statement

The interactome can be requested from the corresponding author. All other data is contained within the article and supplementary files.

## Acknowledgments

The authors would like to thank Dr. Feixiong Cheng and Prof. Albert-László Barabási for providing the current version of the human interactome.

